# Adapting genotyping-by-sequencing and variant calling for heterogeneous stock rats

**DOI:** 10.1101/523043

**Authors:** Alexander F. Gileta, Jianjun Gao, Apurva S. Chitre, Hannah V. Bimschleger, Celine L. St. Pierre, Shyam Gopalakrishnan, Abraham A. Palmer

## Abstract

The heterogeneous stock (**HS**) is an outbred rat population derived from eight inbred rat strains. HS rats are ideally suited for genome wide association studies; however, only a few genotyping microarrays have ever been designed for rats and none of them are currently in production. To address the need for an efficient and cost effective method of genotyping HS rats, we have adapted genotype-by-sequencing (**GBS**) to obtain genotype information at large numbers of single nucleotide polymorphisms (**SNPs**). In this paper, we have outlined the laboratory and computational steps we took to optimize double digest genotype-by-sequencing (**ddGBS**) for use in rats. We also evaluate multiple existing computational tools and explain the workflow we have used to call and impute over 3.7 million SNPs. We also compared various rat genetic maps, which are necessary for imputation, including a recently developed map specific to the HS. Using our approach, we obtained concordance rates of 99% with data obtained using data from a genotyping array. The principles and computational pipeline that we describe could easily be adapted for use in other species for which reliable reference genome sets are available.

## INTRODUCTION

Advances in next-generation sequencing technology over the past decade have enabled the discovery of high-density, genome-wide single nucleotide polymorphisms (**SNPs**) in model systems. Comprehensive assays of the standing genetic variation in these organisms has allowed for the identification of quantitative trait loci (**QTL**) and the application of numerous population genetic and phylogenetic methods. However, due to the high degree of linkage disequilibrium (**LD**) in the populations, sequencing whole genomes is not necessary. Many populations are the result of numerous generations of interbreeding inbred strains, allowing for recombination to produce an admixed population with known founder haplotypes. Due to the relatively slow rate of accumulation of recombination events, these populations contain large chunks of the genome derived from the same founder haplotype. Nearby SNPs are therefore often in strong LD with physically adjacent loci, effectively ‘tagging’ nearby variation and thereby reducing the number of sites that need to be directly genotyped. Several reduced-representation sequencing approaches that take advantage of LD structure have been previously described (Miller et al. 2007; van Orsouw et al. 2007; Van Tassell et al. 2008; Baird et al. 2008; X. Huang et al. 2009; Andolfatto et al. 2011; Davey et al. 2011; Elshire et al. 2011; Poland et al. 2012; Peterson et al. 2012; Sun et al. 2013; Scheben, Batley, and Edwards 2017). Thousands of SNPs can be identified in large numbers of samples for a fraction of the price of whole-genome sequencing methods (Chen et al. 2013; He et al. 2014). The advantages of these methods are especially attractive when applied to less commonly utilized species or strains for which genotyping microarrays are not available.

Of the existing reduced-representation protocols, the genotyping-by-sequencing (**GBS**) approach developed by Elshire et al. (Elshire et al. 2011) has been frequently modified to accommodate other species: soybean (Sonah et al. 2013), rice (Furuta et al. 2017), oat (Fu and Yang 2017), chicken (Pértille et al. 2016; Wang et al. 2017), mouse (Parker et al. 2016), fox (Johnson et al. 2015), and cattle (De Donato et al. 2013), among others. The greatly varying genomic composition among organisms necessitates a diverse and customized set of approaches for obtaining high-quality genotypes. As such, both the GBS protocol and computational pipeline require modifications when applied to a new species. Recent work from our group showed that GBS can be effectively applied to outbred mice (Parker et al. 2016; Gonzales et al. 2017; Zhou et al. 2018) and rats (Fitzpatrick et al. 2013). However, those publications used protocols that had not been optimized, leaving significant room for improvement in genotype quality and marker density. Additionally, although several tools and workflows for the analysis of GBS data have been described, including Stacks (Catchen et al. 2013), IGST-GBS (Sonah et al. 2013), TASSEL-GBS (Glaubitz et al. 2014), Fast-GBS (Torkamaneh et al. 2017), and GB-eaSy (Wickland et al. 2017), the majority were developed and optimized for use in plant species and given the lack of well-developed genomic resources in these species, do not leverage the wealth of genomic data available for model organisms such as rats. Here we describe the customized computational and laboratory protocols for applying GBS to HS rats.

The HS is an outbred rat population created in 1984 using eight inbred strains and has been maintained since then with the goal of minimizing inbreeding and maximizing the genetic diversity of the colony (Johannesson et al. 2008; Woods and Mott 2017). After more than 80 generations of accumulated recombination events, their genome has become a fine-scale mosaic of the inbred founders’ haplotypes. The breeding scheme and the number of accumulated generations has made the HS colony attractive for genetic studies. Additionally, extensive deep sequencing data exists for many inbred rat strains, including the eight progenitor strains (Rat Genome Sequencing and Mapping Consortium et al. 2013; Hermsen et al. 2015; Ramdas et al. 2019), allowing for accurate imputation to millions of additional SNPs.

Detailed here are the steps we have taken to optimize a rat GBS protocol and computational pipeline. Drawing on existing protocols (Elshire et al. 2011; Peterson et al. 2012; Poland et al. 2012; Parker et al. 2016) as templates, we redesigned our previous GBS approach (Parker et al. 2016; Gonzales et al. 2018) and have developed a novel, reference-based, high-throughput workflow to accurately and cost-effectively call and impute variants from low-coverage double digest GBS (**ddGBS**) data in HS rats. This publication is intended as a resource for others who might wish to perform GBS in rats and should provide a roadmap for adapting GBS for use in new species. We demonstrate that with a suitable reference panel, applying reduced representation approaches and imputation in model systems can provide high-confidence genotypes on millions of genome-wide markers.

## MATERIALS AND METHODS

### Tissue samples and DNA extraction

Samples for this study originated from three sources: an in house advanced intercross line (**AIL**) derived from LG/J and SM/J mice (Gonzales et al. 2018), Sprague Dawley (**SD**) rats from Charles River Laboratories and Harlan Sprague Dawley, Inc. (Gileta et al. 2018), and an HS rat colony (Woods and Mott 2017; Chitre et al. 2018). Early stages of ddGBS optimization utilized AIL genomic DNA extracted from spleen by a standard salting-out protocol. Later optimization steps were performed using genomic DNA from SD rats extracted from tail tissue using the PureLink Genomic DNA Mini Kit (Thermo Fisher Scientific, Waltham, MA). HS rat DNA was extracted from spleen tissue using the Agencourt DNAdvance Kit (Beckman Coulter Life Sciences, Indianapolis, IN). All genomic DNA quality and purity was assessed by NanoDrop 8000 (Thermo Fisher Scientific, Waltham, MA). Interestingly, we observed that rat genomic DNA derived from either spleen or tail tissue appears to degrade faster than mouse genomic DNA following extraction by either of the above protocols; therefore, we recommend storing rat genomic DNA at −20° and using it within weeks of extraction whenever possible.

### *In silico* digest of rat genome

We used *in silico* digests to aid in the selection of restriction enzymes, with the goal of maximizing the proportion of the genome captured at sufficient depth to make confident genotype calls (Kent et al. 2002). We used the *restrict* function in EMBOSS (version 6.6.0) (Rice, Longden, and Bleasby 2000) in conjunction with the REBASE database published by New England BioLabs (NEB; version 808) (Roberts and Macelis 1999) to perform *in silico* digest of the current release of the Norway brown rat reference genome, designated rn6. For the primary restriction enzyme, we chose PstI, which had been successfully used in numerous project (Fitzpatrick et al. 2013; Parker et al. 2016; Gonzales et al. 2018). We performed the digest with PstI alone and then with PstI paired with each of 7 secondary enzymes: AluI, BfaI, DpnI, HaeIII, MluCI, MspI, and NlaIII. We only considered fragments with one PstI cut site and one cut site from the secondary enzyme because the adapter and primer sets are designed to only allow these fragments to be amplified.

### Restriction enzyme selection

Initial criteria for selecting a secondary restriction enzyme were a 4bp recognition sequence, no ambiguity in the recognition sequence (i.e. N’s), compatibility with the NEB CutSmart Buffer, and an incubation temperature of 37°C. The list of enzymes meeting these criteria at the time included AluI, BfaI, DpnI, HaeIII, MluCI, MspI, and NlaIII. Using the *in silico* digest data, we looked to maximize the portion of the genome contained within a fragment size range of 125-275bp (250-400bp with annealed adapters and primers) (Figure 1; Table 1). We excluded enzymes that produced blunt ends, both because it would be more difficult to anneal adapters to blunt ended fragments and because our adapters would then also anneal to blunt ends produced by DNA shearing. We also excluded methylation-sensitive enzymes, as we did not want to limit the breadth of our sequencing efforts, accepting the possibility of read pileup in repetitive regions. Based on these criteria, as well as maximizing the percent of the genome captured, NlaIII, BfaI, and MluCI were selected for further testing. The final choice of enzyme (NlaIII) was determined empirically and is detailed in the Results.

**Table 1.** Restriction enzyme options for double digest. The percent genome in region columns indicate the percentage of the genome that falls within the provided fragment size ranges and can therefore be captured by GBS.

| Restriction Enzyme(s) | Recognition sequence | Length of Overhang (bp) | % Genome in 250-400bp Region<sup>+</sup> | % Genome in 300-450bp Region<sup>+</sup> |
| --- | --- | --- | --- | --- |
| PstI | CTGCA<sup>^</sup>G | 4 | 0.48% | 0.56% |
| PstI + AluI | AG<sup>^</sup>CT | 0 | 3.06% | 2.88% |
| PstI + BfaI | C<sup>^</sup>TAG | 2 | 3.10% | 3.25% |
| PstI + DpnI* | GA<sup>^</sup>TC | 0 | 2.69% | 3.00% |
| PstI + HaeIII | GG<sup>^</sup>CC | 0 | 2.71% | 2.79% |
| PstI + MluCI | <sup>^</sup>AATT | 4 | 3.32% | 3.21% |
| PstI + MspI | C<sup>^</sup>CGG | 2 | 1.16% | 1.24% |
| PstI + NlaIII | CATG<sup>^</sup> | 4 | 3.45% | 3.31% |

\* Restriction enzyme is methylation sensitive.
<sup>+</sup> Calculated using rn6 genome length of 2,870,182,909bps.

**Figure 1.**
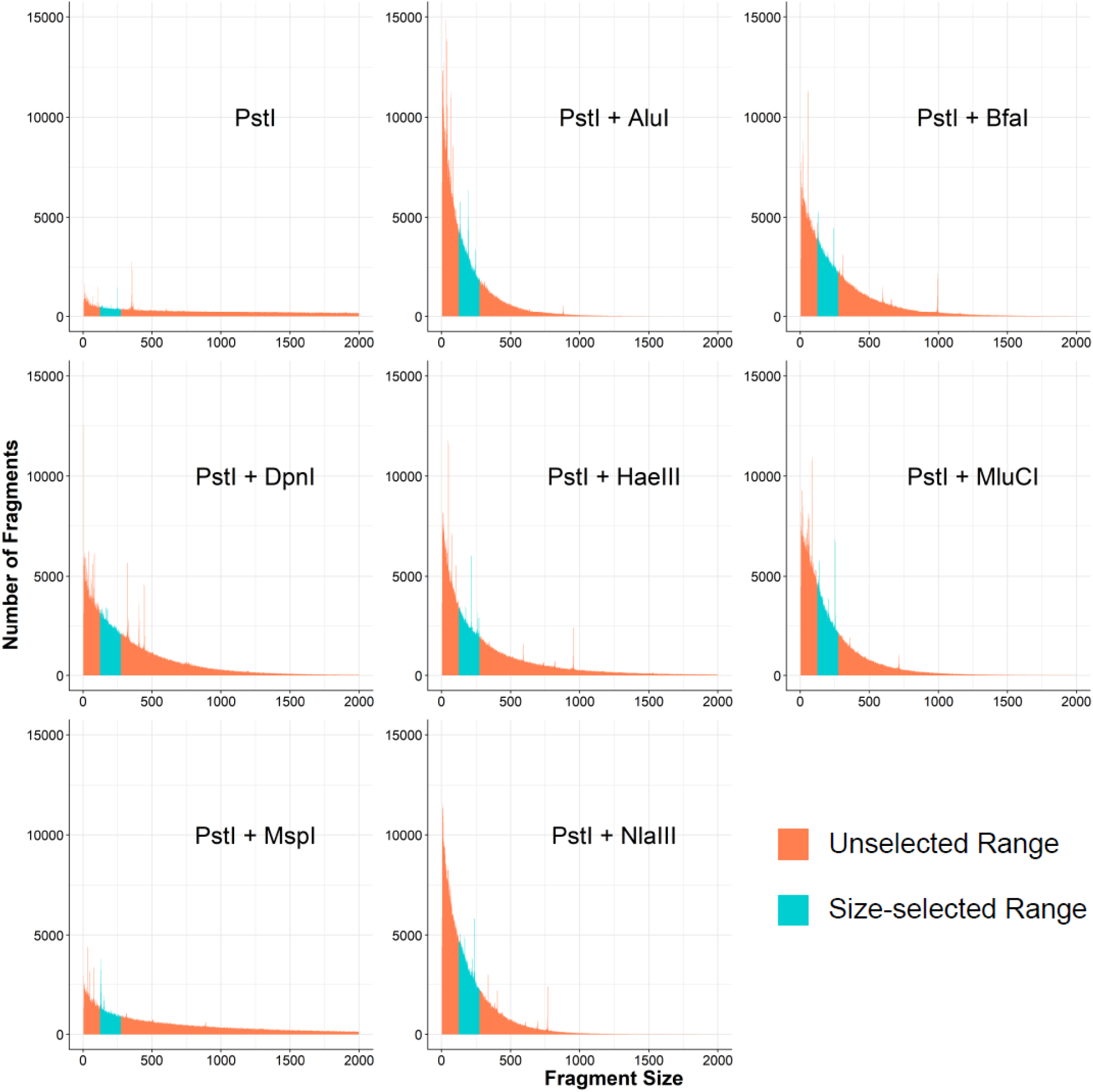
*In silico* digest fragment distributions for PstI and potential secondary restriction enzymes. Each panel represents an independent digest of rn6 with the listed enzyme(s). Regions highlighted in blue are fragments that would be selected by the Pippin Prep (125-275bp) after annealing adapters and primers. These regions are quantified in Table 1 by multiplying the length of the fragments by the number of fragments to estimate the portion of the genome captured.

### ddGBS library preparation and sequencing

The full ddGBS protocol is available in File S1. In brief, approximately 1μg of DNA was used per sample. Sample DNA, PstI barcoded adapters, and NlaIII Y-adapter were combined in a 96-well plate and allowed to evaporate at 37°C overnight. The PstI adapter barcode is “in-line” such that each sequencing read from a given sample with contain both the PstI overhang sequence (4bps) and unique adapter sequence (4-8bps) prior to the beginning of the insert sequence. Sample DNA and adapters were re-eluted on day two with a PstI/NlaIII digestion mix and incubated at 37°C for two hours to allow for complete digestion. Ligation reagents were then added and incubated at 16°C for one hour to anneal the adapters to the DNA fragments, followed by a 30-minute incubation at 80°C to inactivate the restriction enzymes. Sample libraries were purified using a plate from a MinElute 96 UF PCR Purification Kit (QIAGEN Inc., Hilden, Germany), vacuum manifold, and ddH_2_O. Once re-eluted, libraries were quantified in duplicate with Quant-IT PicoGreen (Thermo Fisher Scientific, Waltham, MA) and pooled to the desired level of multiplexing (i.e. 12, 24, or 48 samples per library). Each pooled library was then concentrated by splitting the pooled volume across 2-3 wells of the MinElute vacuum plate and resuspending the library at desired volume for use in the Pippin Prep. The concentrated pool was quantified to ensure the gel cassette will not be overloaded with DNA (>5μg). The pool was then loaded into the Pippin Prep for size selection (300-450bps) using a 2% agarose gel cassette on a Pippin Prep (Sage Science, Beverly, MA). Size-selected libraries were then PCR amplified for 12 cycles to add the Illumina sequencing primers and increase the quantity of DNA, concentrated again, and checked for quality on an Agilent 2100 Bioanalyzer with a DNA 1000 Series II chip (Agilent Technologies, Santa Clara, CA)., Bioanalyzer results were used to assure sufficient DNA concentration and to identify excessive primer dimer peaks.

As a pilot, an initial 96 HS samples were sequenced, 12 samples per library, at Beckman Coulter Genomics (now GENEWIZ) on an Illumina HiSeq 2500 with v4 chemistry and 125bp single-end reads. Subsequently, we began using a set of 48 unique barcoded adapters (File S2) to multiplex 48 HS samples per ddGBS library. Each library thereafter was run on a single flow cell lane on an Illumina HiSeq 4000 with 100bp single-end reads at the IGM Genomics Center (University of California San Diego, La Jolla, CA). We also obtained ddGBS data on the HiSeq 4000 for a select set of 96 samples that had been previously genotyped on a custom Affymetrix Axiom MiRat 625k microarray (Part#: 550572), providing us with a truth set.

### Evaluation of ddGBS pipeline performance

We present the steps required to call and impute genotypes from raw ddGBS sequencing data in Figure 2. During optimization of the pipeline, performance was assessed by two primary metrics: (1) the number of variants called and (2) genotype concordance rates for calls made in 96 HS rats that had both ddGBS genotypes and array genotypes from a custom Affymetrix Axiom MiRat 625k microarray. There were two checkpoints in the GBS pipeline where genotype quality (measured by concordance rate) was assessed. The first was after “internal” imputation with Beagle (Browning and Browning 2009; 2016), whereby we leverage information from samples that had sufficient read depth to make a confident genotype call at a given locus in order to impute the genotype of other samples that had lower read depths at that locus. The second checkpoint was after “external” imputation, meaning imputation to our reference panel with IMPUTE2 to obtain genotype calls at loci we did not directly capture by our GBS method. (B. N. Howie, Donnelly, and Marchini 2009; B. Howie et al. 2012). A third, additional metric we checked was the transition to transversion ratio (TSTV), which is expected to be ~2 for intergenic regions. The steps as outlined in the following sections reflect the final version of the pipeline. Variant calling and imputation steps utilized all available samples run on the HiSeq 4000 (3,000+ rats), though genotype concordance rates could only be calculated for the set of 96 HS samples for which we had array genotype calls.

**Figure 2.**
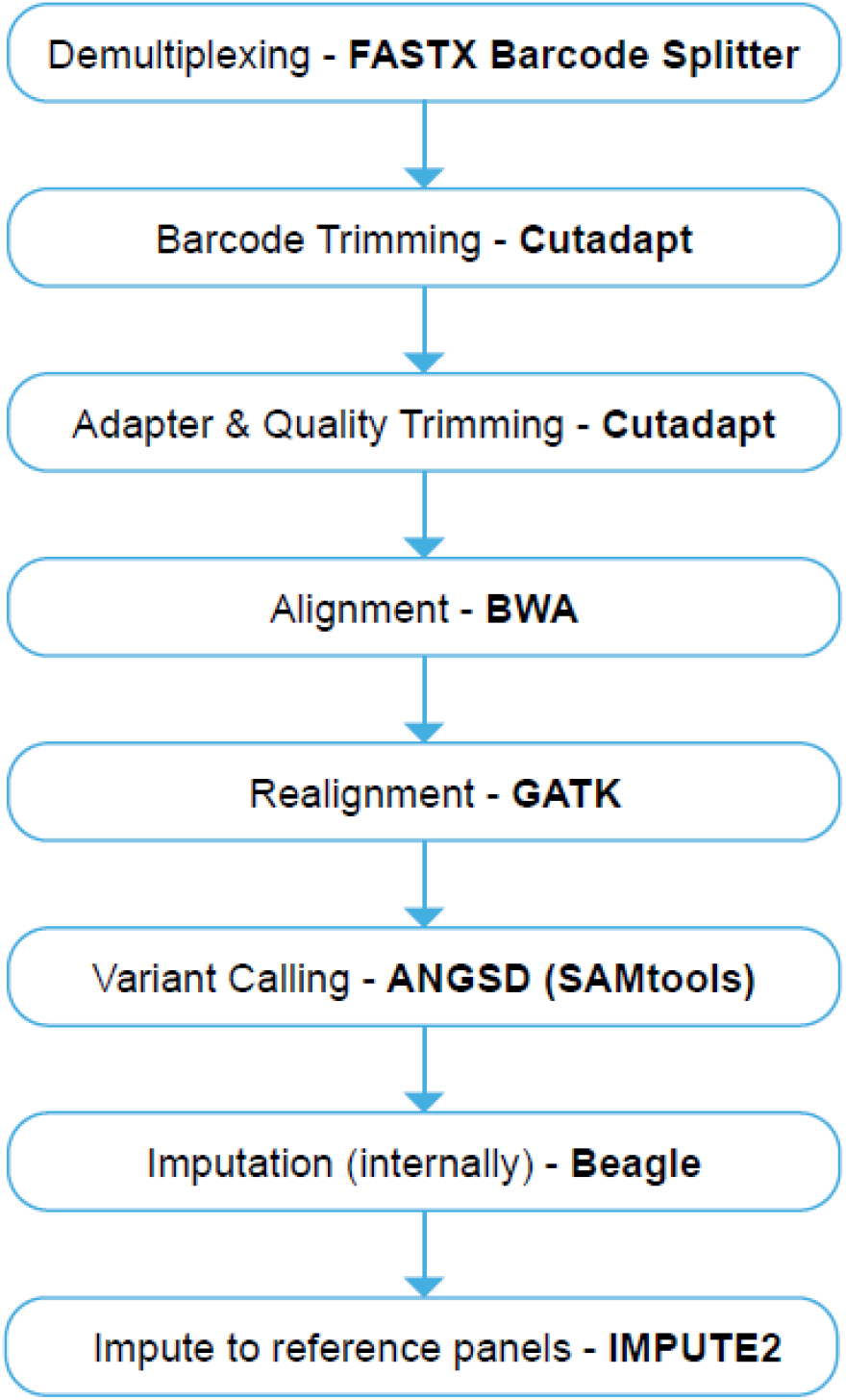
ddGBS sequencing data analysis workflow. Each step of the workflow is described in the text.

### Demultiplexing

The PstI adapter barcodes were used to demultiplex FASTQ files into individual sample files. Three demultiplexing software packages were tested: FASTX Barcode Splitter v0.0.13 [RRID: SCR_005534] (Hannon Lab 2010), GBSX v1.3 (Herten et al. 2015), and an in-house Python script (Parker et al. 2016). Reads that could not be matched with any barcode (maximum of 1 mismatch allowed), or that lacked the appropriate enzyme cut site, were discarded. Samples with less than two million reads after demultiplexing were discarded as these appeared to be outliers (Figure S4) and were observed to have high rates of missingness in their genotype calls. Data concerning demultiplexing are shown in Table S1 are from a single HS rat sequenced in a 12-sample library on one lane after demultiplexing and adapter/quality trimming.

### Barcode, adapter, and quality trimming

Read quality was assessed using FastQC v0.11.6 (Andrews 2017). We compared the efficacy of two rapid, lightweight software options for trimming barcodes, adapters, and low-quality bases from the NGS reads: Cutadapt v1.9.1 (Martin 2011) and the FASTX Clipper/Trimmer/Quality Trimmer tools v0.0.13 (Hannon Lab 2010) (Table S2). A base quality threshold of 20 was used and reads shorter than 25bp were discarded.

### Read alignment and indel realignment

Rattus norvegicus genome assembly rn6 was used as the reference genome for read alignment with the Burrows-Wheeler Aligner v0.7.5a (BWA) [RRID: SCR_010910] (H. Li and Durbin 2009) using the *mem* algorithm. We then used GATK IndelRealginer v3.5 [RRID: SCR001876] (McKenna et al. 2010) to improve alignment quality by locally realigning reads around a reference set of known indels in 42 whole-genome sequenced inbred rat strains, including the eight HS progenitor strains (Hermsen et al. 2015).

### Variant calling

Variants were called, and genotype likelihoods were computed at variant sites using ANGSD v0.911, under the SAMtools model for genotype likelihoods (ANGSD-SAMtools) (Korneliussen, Albrechtsen, and Nielsen 2014; Durvasula et al. 2016). Further, using ANGSD-SAMtools, we inferred the major and minor alleles (*-domajorminor* 1) from the genotype likelihoods, retaining only high confidence polymorphic sites (*-snp_pval* 1e-6), and estimated the allele frequencies based on the inferred alleles *(-domaf* 1). We discarded sites missing read data in more than 4% of samples (*-minInd*). Additionally, we tested multiple thresholds for minimum base (*-minQ*) and mapping (*-minMapQ*) qualities.

### Internal imputation

Beagle v4.1 (Browning and Browning 2009; 2016) was used to improve the genotyping within the samples without the use of an external reference panel. Missing and low quality genotypes were imputed by borrowing information from other individuals in the dataset with high quality information at these same variant sites. Before settling on the combination of ANGSD-SAMtools and Beagle for genotype calling and internal imputation, we also experimented with GATK’s HaplotypeCaller (McKenna et al. 2010) with various parameter settings, but with unsatisfactory results (Figure 3).

**Figure 3.**
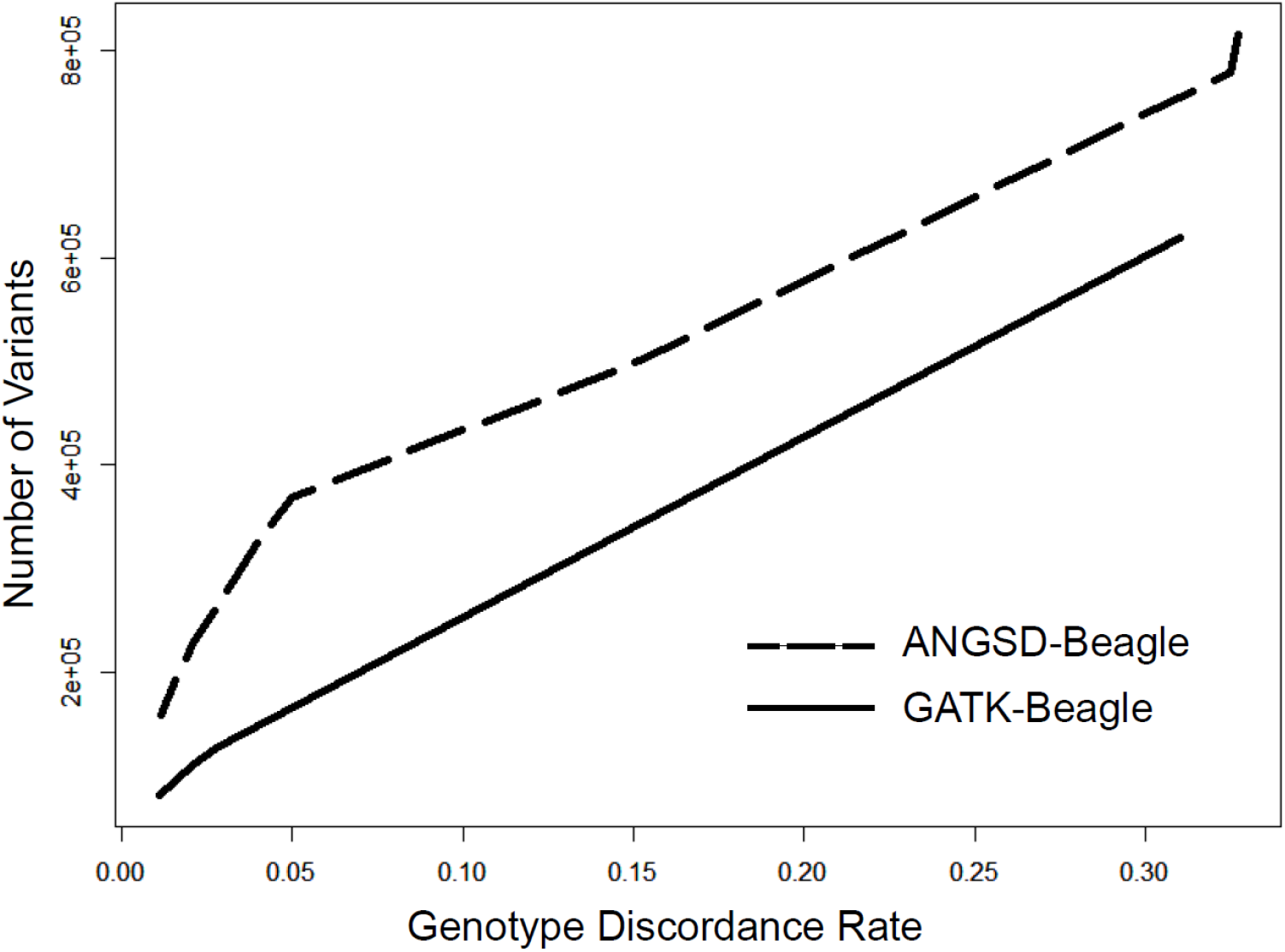
Genotype discordance rates between array data and variants called by GATK/Beagle or ANGSD-SAMtools/Beagle. The figure compares the number variants called by combination of ANGSD-SAMtools and Beagle or GATK HaplotypeCaller and Beagle at various thresholds of genotype discordance with array data. Calls were made using the 96 HS rats with array data. The x-axis represents the genotype discordance rate thresholds and the y-axis is the number of variants that surpass that threshold for each genotype calling method.

### Quality Control for genotypes before imputation using an external reference panel

To verify the quality of the “internally” imputed genotypes prior to imputing SNPs from the 42 inbred strain reference panel (Hermsen et al. 2015), we checked concordance rates for the 96 HS animals with array genotypes, examined the TSTV ratio, and assessed whether the sex as recorded in the pedigree records agreed with the sex empirically determined by the proportion of reads on the X-chromosome out of the total number of reads (Figure S1). We also identified Mendelian errors using the --mendel option in *plink* and known pedigree information for 1,136 trios from 214 families within the HS sample. Using the fraction of the trios that were informative for a given SNP and the formula 1-(1-2p(1-p))^3^, where p represents the minor allele frequency of the allele, we formed curves for the distributions of the expected number of Mendelian errors for both SNPs and samples and chose the inflection points as thresholds for the number of Mendelian errors allowed.

### Data preparation for phasing with external reference panel

First, in our study sample of 96 samples, we only retained variants previously identified in the 8 HS founder strains because we expected the polymorphisms in our samples to be limited to the variation present in the founders (Hermsen et al. 2015; Ramdas et al. 2019). Further, to improve imputation efficiency, we employed a pre-phasing step with IMPUTE2 (*prephase_g*) (B. Howie et al. 2012) prior to imputation. Pre-phasing only needs to be performed once, allowing us to reuse the estimated haplotypes from out dataset for imputation with multiple different reference panels. A flowchart outlining the pre-phasing protocol is presented in Figure S2.

### Genetic maps

Genetic maps are required for phasing and imputation with IMPUTE2. When we began this project, no strain-specific recombination map was available for HS rats. Thus, we considered a sparse genetic map for SHRSPxBN (Steen et al. 1999). We also tested two types of linearly interpolated genetic maps, with recombination rates set at either 1cM/Mb or the chromosome specific averages for rats, as reported by Jensen-Seaman et al. (Jensen-Seaman 2004). Lastly, late in the course of this project, we experimented with an HS-specific genetic map developed by Littrell et al. (Littrell et al. 2018).

### Imputation to reference panel

We used a combination of existing sequencing and array genotyping data from the HS rat founder and other inbred laboratory rat strains (Hermsen et al. 2015) as reference panel for imputation. Genotype data underwent QC and were phased by Beagle into single chromosome haplotype files. Haplotype files were then created using the workflow detailed in Figure S2. Imputation by IMPUTE2 was performed in 5Mb windows using the aforementioned reference panels and genetic maps.

### Data availability

Genotype data will be available at https://dx.doi.org/10.6084/m9.figshare.8243222. The code necessary to run the steps outlined in the publication are provided at https://dx.doi.org/10.6084/m9.figshare.8243156. Supplementary Files are available at https://dx.doi.org/10.6084/m9.figshare.8243129. Remaining files necessary for imputation (genetic maps, reference data, etc.) can be found with the following links: https://dx.doi.org/10.6084/m9.figshare.11919615, https://dx.doi.org/10.6084/m9.figshare.11919573, https://dx.doi.org/10.6084/m9.figshare.11919597.

## RESULTS

### ddGBS optimization

Previous projects utilizing GBS in mice and rats (Fitzpatrick et al. 2013; Parker et al. 2016; Gonzales et al. 2018) often encountered an issue where certain regions of the genome experienced high pileups of reads per sample (>100x), while other regions were covered by just 1-2 reads. This read distribution imbalance can be caused in part by PCR amplification bias, where a subset of fragments are preferentially amplified until they dominate the final library (Kanagawa 2003; Aird et al. 2011). These previous protocols utilized 18 cycles of amplification. We tested reducing this to 8, 10, 12, or 14 cycles and found that below 12 cycles, there was insufficient PCR product to accurately quantify and pool for sequencing. The reduction in the number of PCR cycles was expected to reduce PCR bias, though this was not explicitly tested.

Another concern regarding previous sequencing results was an excess of long fragments (>700bps as determined by *in silico* digest), which do not provide sufficient reads to make confident genotype calls (< 5 reads per sample) and are therefore wasteful. We tested three methods of combating this issue, including increasing the digestion time or enzyme concentration, performing size selection on the libraries, and using a two-enzyme restriction digest.

We considered the possibility that the restriction enzyme digests might not be running to completion. To address this possibility, we increased the duration of the digestion from 2 hours to 3 or 4 hours. We also tried increasing the number of units of PstI enzyme added, to ensure complete digest. Neither of these modifications impacted the final fragment length distribution of the library, indicating that the digest was reaching completion after 2 hours using the original concentration of PstI (File S3 - wells 1-6).

Our previous GBS protocol did not have an explicit library fragment size selection step. The final library was purified using a MinElute PCR Purification Kit (QIAGEN Inc., Hilden, Germany), which isolates PCR products 70bp-4kb in length, leaving a wide range of fragment sizes in the final library, under the assumption that only shorter fragments would bridge amplify on the flow cell. This method was imprecise and had low reproducibility, negatively impacting our ability obtain reads at consistent sites across libraries. Rather than attempt size selection by gel extraction, we chose to utilize a Pippin Prep, which automates the elution of DNA libraries of desired fragment size ranges. By using this automated size selection, we reduced the proportion of the genome targeted for sequencing, Additionally, since restriction enzymes make predominantly consistent cuts across samples (barring the presence of polymorphisms in RE recognition sites), it is ensured that highly similar sets of genomic fragments will be sequenced across sample libraries. Since the clustering process involves a bridge amplification step that preferentially amplifies library fragments with shorter insert sizes (Illumina, Inc. 2014), we kept the size selection window narrow (250-400bps) to avoid introducing a bias in which fragments were sequenced. A comparison of the fragment size distributions for the protocols before and after introduction of the Pippin Prep is shown in File S4.

To increase the proportion of the genome captured within the fragment size window, we pursued a double digest of the genome using a secondary enzyme with a more frequently occurring recognition sequence. When used alone, *in silico* digest of the rn6 reference genome by PstI (Figure 1; Table 1) showed that only ~0.5% of the genome would have fallen within a 150bp fragment size window selected on the Pippin Prep. Previously, we performed GBS in CFW mice using the single-enzyme approach and observed that large regions of the genome that were not covered by sequencing reads (Parker et al. 2016). Therefore, we sought to increase the fraction of the genome that was accessible to GBS, so that there would be sufficient SNPs to tag the majority of the variation in the rat genome. Additionally, we were concerned about potential biases in coverage, heterozygosity, and the minor allele frequency (**MAF**) spectrum that may be introduced by a less complete capture of the genome. Flanagan and Jones have performed an empirical study comparing single- to double- digest RAD-seq and found that double-digest RAD-seq had lower rates of allelic dropout, decreased variance in between-sample per SNP coverage, less allele frequency inflation due to PCR bias, and reduced batch effects (Flanagan and Jones 2018).

The number of fragments with one of each of the cut sites were summed for all observed lengths and the results summarized in Figure 1 and Table 1. BfaI, MluCI, and NlaIII were chosen for further testing due to their compatibility with PstI digestion reagents and temperatures, sticky ends, and the proportion of the genome falling in the size selection window. We ruled out BfaI because it only had a 2bp overhang after cleavage, which led to a high concentration of adapter dimer in the sequencing libraries (S5 File). NlaIII was chosen because it contained the greatest portion of the genome in the size selection window.

In our previous GBS protocol, all fragments were cut on both ends by PstI. By using a substantially lower concentration of the barcoded PstI adapter than the common PstI adapter, we ensured the barcoded adapter would be the limiting reagent and the majority of fragments with an annealed barcoded adapter would have a common adapter on the other end. This is crucial, as having one of each of the adapters is required for proper amplification of the fragments on the flow cell. However, when using both PstI and NlaIII, the library is predominantly composed of fragments cut on both sides by NlaIII (File S6), which will amplify during PCR with a common adapter, but not on the flow cell. Therefore, we employed a Y-adapter (Poland et al. 2012) to control the direction of the first round of PCR and prevent two-sided NlaIII fragments from dominating the final sequencing library (File S2).

We tested numerous quantities of PstI and NlaIII adapters in an attempt minimize the amount used and avoid adapter dimers in the final libraries. For the barcoded PstI adapters, we tested 120pmol, 60pmol, 20pmol, 4.0pmol, 2.67pmol, 1.60pmol, 0.53pmol, and 0.20pmol; for the NlaIII Y-adapter, 30pmol, 10pmol, 5.0pmol, 4.0pmol, and 1.0pmol (Files S7 & S8). We found that 0.20pmol of PstI adapter and 4pmol of NlaIII Y-adapter yielded sufficient library and minimized the presence of adapter dimers.

We sequenced a trial flow cell with 8 pooled ddGBS libraries of 12 SD rat samples each (96 total) on a HiSeq 2500 (Illumina, San Diego, CA) with 125bp reads and v3 chemistry, obtaining an average of 15.3 million reads per sample. Given the NlaIII *in silico* digest results suggested we were capturing ~3.4% of the genome and that we were using 125bp reads, this corresponded to approximately 20x coverage of captured sites. We subsequently increased the number of samples to 48 per library for the HS rats because we hypothesized 5x would be sufficient coverage per sample when utilizing imputation to a reference panel. We also discovered that a portion of the reads contained sequence fragments of the NlaIII adapter sequence, indicating there were fragments with insert sizes smaller than 125bps in the final library. To avoid this, we increased the fragment size range to 300-450bps (Table 1), which corresponds to a 175-325bp insert size once the adapters and primers are accounted for. We noted however that the library size distribution obtained from the Pippin Prep was uniformly shifted towards higher fragment lengths (Figure S3). This is a result of the high concentrations of our libraries after pooling and loading the gel cartridge near the upper limit of the recommend number of micrograms of DNA, which can cause slower migration of the DNA across the gel matrix.

The final ddGBS protocol can be found in File S1 and the necessary primer and adapter sequences in File S2. This protocol was used for the sequencing of all HS rats included in the following computational optimization steps. These steps were optimized using

### Demultiplexing

The number of base pairs of sequencing data retained after demultiplexing was fairly consistent across demultiplexing software (Table S1). We ultimately decided to use FASTX Barcode Splitter because it yielded the greatest number of reads after quality/adapter trimming and had faster run times. An average of 330 million 100bp reads were obtained per library, resulting in ~7 million reads per sample. Figure S4 shows the distribution of reads counts for all samples after demultiplexing.

### Adapter and quality trimming

Read quality was substantially improved after trimming the barcode and adapter sequences and low-quality base pairs at the ends of reads (Figure S5). Overall read counts were only marginally reduced by quality trimming (Table S1). We observed that the number of called variant sites and the genotyping rate were both greater when using reads initially processed by cutadapt (Martin, 2011) than reads processed by the FASTX_Toolkit (Table S2). Importantly, a large portion of the additional identified variants were known variant sites from the 42 inbred strains reference set (Figure S6), indicating the elevated call rate was at least in part due to capturing more true variant sites. We viewed this as sufficient support for proceeding with cutadapt for adapter removal and quality trimming.

### Mapping quality

The number of called variants and genotype call rates were identical at read mapping quality (mapQ) thresholds of either 20 or 30 (Table S3) within ANGSD. As the ANGSD mapQ threshold was raised to 45, there was a small reduction in the number of called variants, and then much greater losses at thresholds of 60 or 90. Fortunately, discordance rates between ddGBS and array genotypes were stable at both low and high mapQ thresholds, despite the putatively higher quality of the alignments (Figure S7). This permitted us to select a lower mapQ threshold (mapQ = 20), maximizing the number of variants called without sacrificing genotyping accuracy.

### Variant calling

Figure 3 shows that across all levels of genotype discordance rates (with the array genotyping data), the combination of the ANGSD-SAMtools with BEAGLE produced more SNPs, at various discordance thresholds, than GATK’s HaplotypeCaller (McKenna et al. 2010; DePristo et al. 2011). This observation held when variants were limited only to biallelic sites and SNPs with an MAF > 0.05 (Figure S8). We speculate that the poorer performance of HaplotypeCaller may be in part to the sparsity and non-uniform distribution of GBS genotype data across the genome and the high level of genotype call missingness across samples prior to imputation.

ANGSD supports four different models for estimating genotype likelihoods: SAMtools, GATK, SOAPsnp and SYK. We compared the methods to determine which produced the most SNPs with the lowest error rates. The SOAPsnp model demonstrated an advantage in genotype accuracy and number of variants called post-imputation with Beagle (Figure S9). However, SOAPsnp requires considerably more time (1.7x for 96 samples) and memory and scales poorly with sample size. With greater than 2,000 samples, we were unable to allocate sufficient memory for the SOAPsnp model to successfully run, even after dividing the chromosomes into several, smaller chunks. The marginal benefits of SOAPsnp in accuracy and number of variants were far outweighed by its limitations when applied to a large sample set. The GATK model showed a greater number of variants for more lenient genotype discordance rate threshold. This is in contrast with what was observed in Figures 3 and Figure S8 because ANGSD utilizes the direct genotype likelihood method of the first implementation of GATK, whereas we had previously tested GATK’s HaplotypeCaller. Interestingly though, as the stringency for discordance rate increased, the number of variants converged across the SAMtools, GATK, and SOAPsnp models. We proceeded with the SAMtools model for genotype likelihood estimation due to its previous support in the GBS literature (Torkamaneh et al. 2017), accepting a nominal decrease in highly concordant variants (Figure S9) for a large reduction in run time and memory usage.

### Imputation to reference panel

Imputation is used in two complimentary ways in our protocol. As described earlier, after ddGBS, not all samples will have sufficient sequencing coverage at captured polymorphic loci to make a confident genotype call. Therefore, we first use imputation from other well-covered samples to “fill in the blanks” and assign genotypes to SNP loci in the subset of individuals that lacked confident calls at these sites. imputation in this section to call genotypes at sites where GBS that were inaccessible to ddGBS sequencing. After these missing genotypes have been imputed in all samples, we then use the genotype information we have for the SNPs captured by ddGBS along with the reference panel data on the original 8 HS founders (Hermsen et al. 2015; Ramdas et al. 2019) to impute genotype calls at sites that were inaccessible to ddGBS sequencing. Thus, our second application of imputation is similar to the human genetics application in which imputation using 1000 Genomes (1000 Genomes Project Consortium et al. 2010) increases the number of SNPs beyond those included on a given microarray platform. IMPUTE2 was selected over Beagle for this application because it is able to handle multiple reference panels, allowing us to provide data from all 8 HS founder strains.

Before starting this imputation step, we observed an inflated transition/transversion ratio (Table S4) in our ANGSD-SAMtools/Beagle SNPs. This issue was ameliorated when the SNP set was filtered for only “known” variants that were previously identified in either the 42 inbred strains (Hermsen et al. 2015) or the 8 deep-sequenced HS founders (Ramdas et al. 2019). For imputation, we therefore only provided IMPUTE2 with previously identified variant sites from our ANGSD-SAMtools/Beagle output. Prior to running IMPUTE2, we also filtered the variants for biallelic sites with a genotype call in at least two individuals. Using pedigree data for the HS rats, we further removed samples showing an order of magnitude higher level of Mendelian error than the sample mean. We further removed SNPs that had an error rate surpassing a threshold of ~0.005 (Figure S10; inflection point). There were 4 samples and 4,179 SNPs removed from subsequent analyses. Lastly, we removed any samples where the X chromosome read ratio was incompatible with their reported sex using hard threshold of 3% of total reads, above indicating female and below indicating male (Figure S1).

There were three major genomic reference datasets available for the HS rats. The first reference set was obtained from Baud et al. (Rat Genome Sequencing and Mapping Consortium et al. 2013) and contained sequence data and genotype calls for the 8 founders of the HS. The second came from Hermsen et al. (Hermsen et al. 2015) and sequence and genotype data on 42 distinct laboratory rats strains and substrains, 8 of which were the founders of the HS from Baud et al., but analyzed alongside a new set of strains. The third reference set came from Ramdas et al. (Ramdas et al. 2019), who independently performed whole-genome sequencing and made genotype calls on the 8 HS founder strains. It was unclear which set of genotypes would provide the best reference for imputation from out ddGBS data, so we tested five different possible subsets of available data (Table 2). From Hermsen et al., we used (1) all 42 inbred strains, (2) only the 34 strains that were not the HS founders, and (3) only the 8 HS founder strains. Then from Baud et al. and Ramdas et al., we tested the 8 HS founders only from each study. The most accurate imputation was observed for the reference set containing only the 8 deep-sequenced HS founder strains (Ramdas et al. 2019); however, imputation to this set had the lowest genotyping rate of all panels. In contrast, using the 42 rat inbred strains displayed a balance of high accuracy and low missingness, leading us to choose this as our reference set. To better understand the role of the 8 founder strains, which were part of the 42 strain reference panel, we created a reference panel that included only the 34 non-HS founder strains. As expected, discordance rates were much higher when only considering non-founders. However, the genotype missingness was lower for the 34 than the 8 founders alone, suggesting a combination of the two was the optimal set.

**Table 2.** Imputation accuracy based on different variant reference panels for IMPUTE2. The table includes five different possible reference panels for imputation. The 42 inbred strains, 34 non-founder inbred strains, and 8 HS founders from the 42 inbred strains all were derived from Hermsen et al. 2015 (Hermsen et al. 2015). The UMich 8 HS founders were obtained from Ramdas et al. 2019 (Ramdas et al. 2019). The final set of 8 HS founder was taken from Baud et al. 2013 (Rat Genome Sequencing and Mapping Consortium et al. 2013).

|  |  | **Chr1** | **Chr2** |
| --- | --- | --- | --- |
| **42 inbred strains** | **Discordance rate** | 0.011 | 0.010 |
|  | **# Variants** | 790,659 | 882,993 |
|  | **Genotyping Rate** | 0.85 | 0.81 |
| **All 34 non-founder inbred strains** | **Discordance rate** | 0.035 | 0.030 |
|  | **# Variants** | 812,550 | 912,749 |
|  | **Genotyping Rate** | 0.84 | 0.80 |
| **8 HS founders only from the 42 inbred strains** | **Discordance rate** | 0.012 | 0.011 |
|  | **# Variants** | 805,424 | 902,061 |
|  | **Genotyping Rate** | 0.57 | 0.53 |
| **UMich 8 HS founders only** | **Discordance rate** | 0.0059 | 0.008 |
|  | **# Variants** | 865,514 | 898,621 |
|  | **Genotyping Rate** | 0.42 | 0.41 |
| **Baud et. al 2013 8 HS founders only** | **Discordance rate** | 0.0095 | 0.0096 |
|  | **# Variants** | 507,909 | 540,844 |
|  | **Genotyping Rate** | 0.43 | 0.40 |

IMPUTE2 requires a genetic map. As described in the methods section, we considered four different genetic maps, two that were empirically derived and two that were linear extrapolations based on the physical map (Figure S11). All genetic map performed similarly (Table S5). Surprisingly, the linear genetic maps performed just as well as the HS-specific map (Littrell et al. 2018). Thus, for simplicity, we chose to use the chromosome-specific values initially published by Jensen-Seaman (Jensen-Seaman 2004).

To obtain our final set of ~3.7 million variants, a final round of variant filtering was performed after imputation to the 42 strain reference panel. We removed SNPs with MAF < 0.005, a post-imputation genotyping rate < 90%, and SNPs that violated HWE with p<1×10^−10^.

## DISCUSSION

The use of microarrays and WGS for genotyping large samples in model organisms remains cost-prohibitive. There is therefore an urgent and wide-spread need for high-performance and economical methods of obtaining genome-wide genotype data. While reduced-representation approaches have been utilized in numerous species of plants and animals, including rodents (Peterson et al. 2012; Fitzpatrick et al. 2013; Parker et al. 2016; Gonzales et al. 2017; Zhou et al. 2018), there has yet to be a published protocol optimized specifically for rats. Prior to sequencing thousands of HS samples with GBS for our mapping efforts, we wanted to ensure we were capturing the greatest possible number of high-quality variants at the lowest possible cost. The protocol we present here is the culmination of careful testing and optimization of each step of the GBS protocol for rats. We have now applied the approach to 4,973 HS rats, as well as 4,608 Sprague Dawley rats (Gileta et al. 2018).

Our previous GBS protocol (Parker *et al*, 2016), which was designed for use with CFW mice, was unsuitable for our current genotyping efforts in HS rats, due to the much higher levels of genetic diversity in the HS population. There are multiple reasons we chose to develop our own computational pipeline for GBS rather than using existing workflows. Foremost, the prominent GBS analysis pipelines were developed and optimized for use in crop species (Sonah et al. 2013; Catchen et al. 2013; Glaubitz et al. 2014; Torkamaneh et al. 2017; Wickland et al. 2017), some of which are polyploid and have differing levels of variation and LD than outbred rodent populations. Additionally, there were elements of each pipeline that did not meet our needs or lacked customizability. For instance, TASSEL-GBS v2 (Glaubitz et al. 2014) trims all reads to 92 base pairs; however, other projects underway in our lab utilized up to 125bp reads, leading to a ~20% reduction in data. TASSEL-GBS also ignores read base quality scores, which are informative in probabilistic frameworks for estimating uncertainty in alignments and variant calls (H. Li, Ruan, and Durbin 2008; DePristo et al. 2011; Nielsen et al. 2011), and uses a naïve binomial likelihood ratio method for calling SNPs. Stacks has previously shown poor performance in demultiplexing (Herten et al. 2015; Torkamaneh et al. 2017) and does not make use of the reference genome for priors when calling SNPs (Catchen et al. 2013). Fast-GBS relies on Platypus (Rimmer et al. 2014) for variant calling (WGS500 Consortium et al. 2014; Torkamaneh et al. 2017), which employs a Bayesian method of constructing candidate haplotypes that works poorly with low-pass sequencing data and does not scale well to large sample sizes (Z. Li, Wang, and Wang 2018). Lastly, none of these pipelines included an imputation step, which is crucial for filling in missing genotypes in GBS data and can provide millions of additional SNPs given an appropriate composite reference panel (B. Howie, Marchini, and Stephens 2011; G.-H. Huang and Tseng 2014).

Though we have not explicitly tested each alternate GBS pipeline for the purposes of this publication, this has been recently done by Wickland et al. (Wickland et al. 2017). Their pipeline GB-eaSy, which ours most closely resembles, was found to be superior by a number of metrics to Stacks, TASSEL-GBS, IGST, and Fast-GBS. Similar to GB-eaSy, our pipeline utilizes a double-digest GBS protocol, aligns reads to the reference genome with *bwa mem*, and uses the SAMtools genotype likelihood model for calling SNPs (H. Li 2011). The combination of bwa mem and SAMtools algorithm was independently shown to have the best performance for calling SNPs from Illumina data (Hwang et al. 2015), further supporting our choice of these programs for read alignment and variant calling. Additionally, using the ANGSD wrapper provided us with the ability to convert the posterior genotype probabilities into genotype dosages for mapping studies (Korneliussen, Albrechtsen, and Nielsen 2014).

A minor difference between GB-eaSy and our pipeline is the use of cutadapt (Martin 2011) rather than GBSX (Herten et al. 2015) for demultiplexing, though both performed equally well (Table S1). The primary improvement is our extension of the pipeline with the implementation of effective internal and reference-based imputation steps using the 42 inbred rat genomes (Hermsen et al. 2015) and 8 deep-sequenced HS founders from UMich (Ramdas et al. 2019). There are two stages of imputation in our pipeline. The first one is accomplished by Beagle and aims to fill in missing genotypes at called variants using information from other samples. This raises the genotype call rate to 100%, but it may also introduce errors due to insufficient information, emphasizing the need for careful filtering steps. The second stage of imputation made use of IMPTUE2 and an external reference panels of variants called from WGS data on the 8 inbred HS founders, as well as 34 additional inbred rat strains. We decided to include the 34 additional strains because of the elevated genotyping rate we observed upon their inclusion in the IMPUTE2 reference panel. We attribute this to the presence of haplotypes that exist in both the 8 the HS founder strains and a subset of the 34 additional strains in this panel. The benefits of using a composite reference panel have been previously noted (Zhang et al. 2013; G.-H. Huang and Tseng 2014); there is increased accuracy and decreased missingness in the imputed genotype data.

In summary, we have adapted a GBS protocol and genotyping and imputation pipeline to obtain dense genotypes on genome-wide markers in highly-multiplexed HS rats. After quality filtering on the level of SNP and sample, over 3.7 million SNPs were called with a concordance rate of 99%. The ddGBS protocol and bioinformatic methods used to produce this data are publicly available, easy to handle, and cost-effective. The presented workflow could be feasibly followed with marginal modifications for application in other species. The steps taken toward optimizing the wet lab protocols are easily applied to novel organisms, as is the computational pipeline so long as there are reliable reference genome sets available for use in alignment and imputation.

## ACKNOWLEDGEMENTS

This work was supported by P50 DA037844, R21 DA036672, T32 GM07197, and F31 DA039638.

---

**Figure S1.**
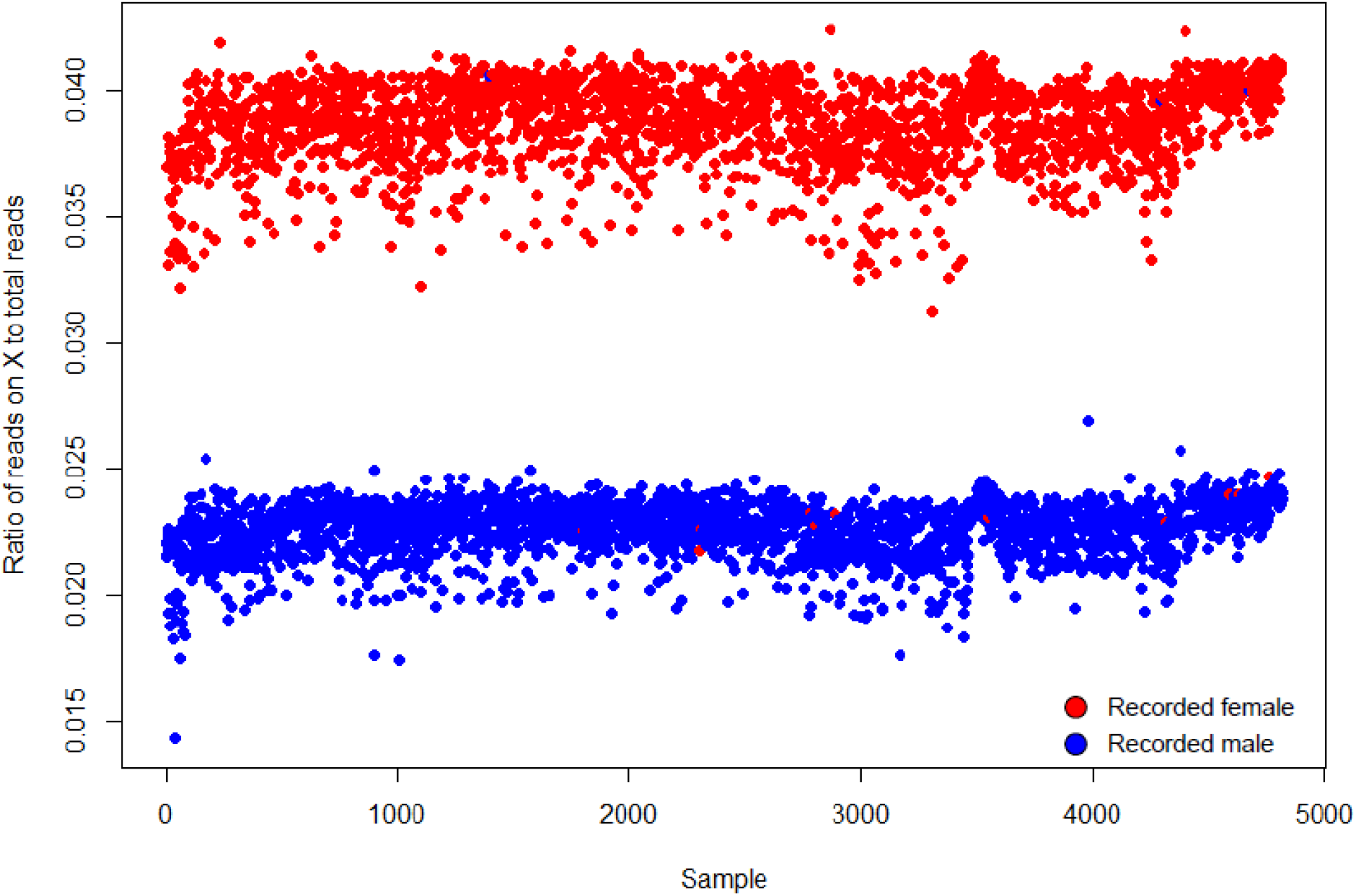
Ratio of reads on X-chromosome to total sequencing reads. The color of the points indicates the pedigree-recorded sex of the samples. Females are expected to have approximately twice as many reads for the X-chromosome. Samples that did not cluster with their pedigree-recorded sex were removed from the study for possible sample mix-up.

**Figure S2.**
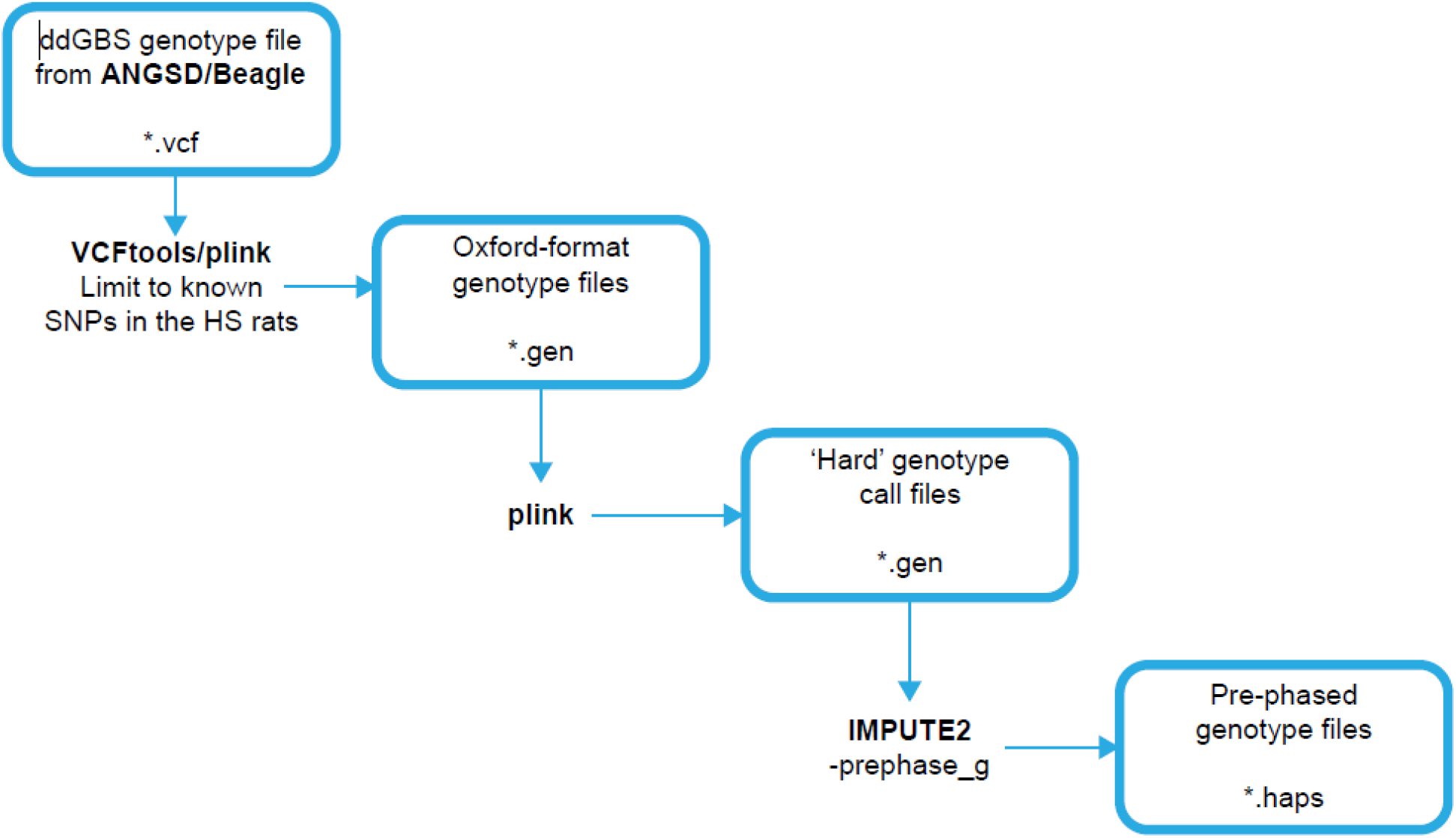
Data preparation workflow for imputation with IMPUTE2.

**Figure S3.**
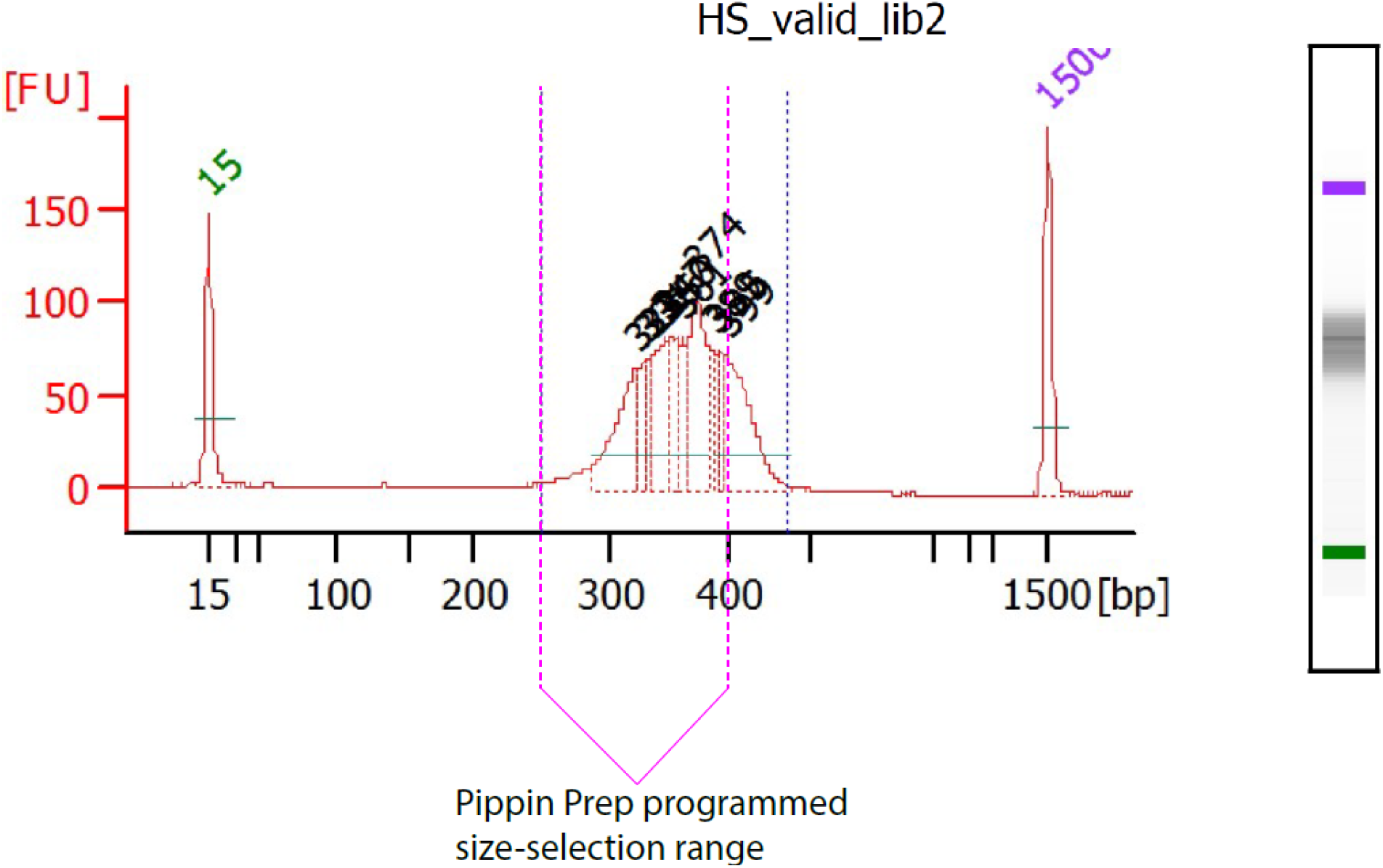
Programmed vs. empirical Pippin Prep fragment size range. This plot comes from the Bioanalyzer output for a pooled HS library. The x-axis shows the library fragment sizes in base pairs, and the y-axis is in fluorescent units, which represent the quantity of the fragments on the gel chip. There is approximately a 50-75bp shift in the empirical library distribution compared to expectation due to the high quantity of fragments loaded into the Pippin Prep gel cassette.

**Figure S4.**
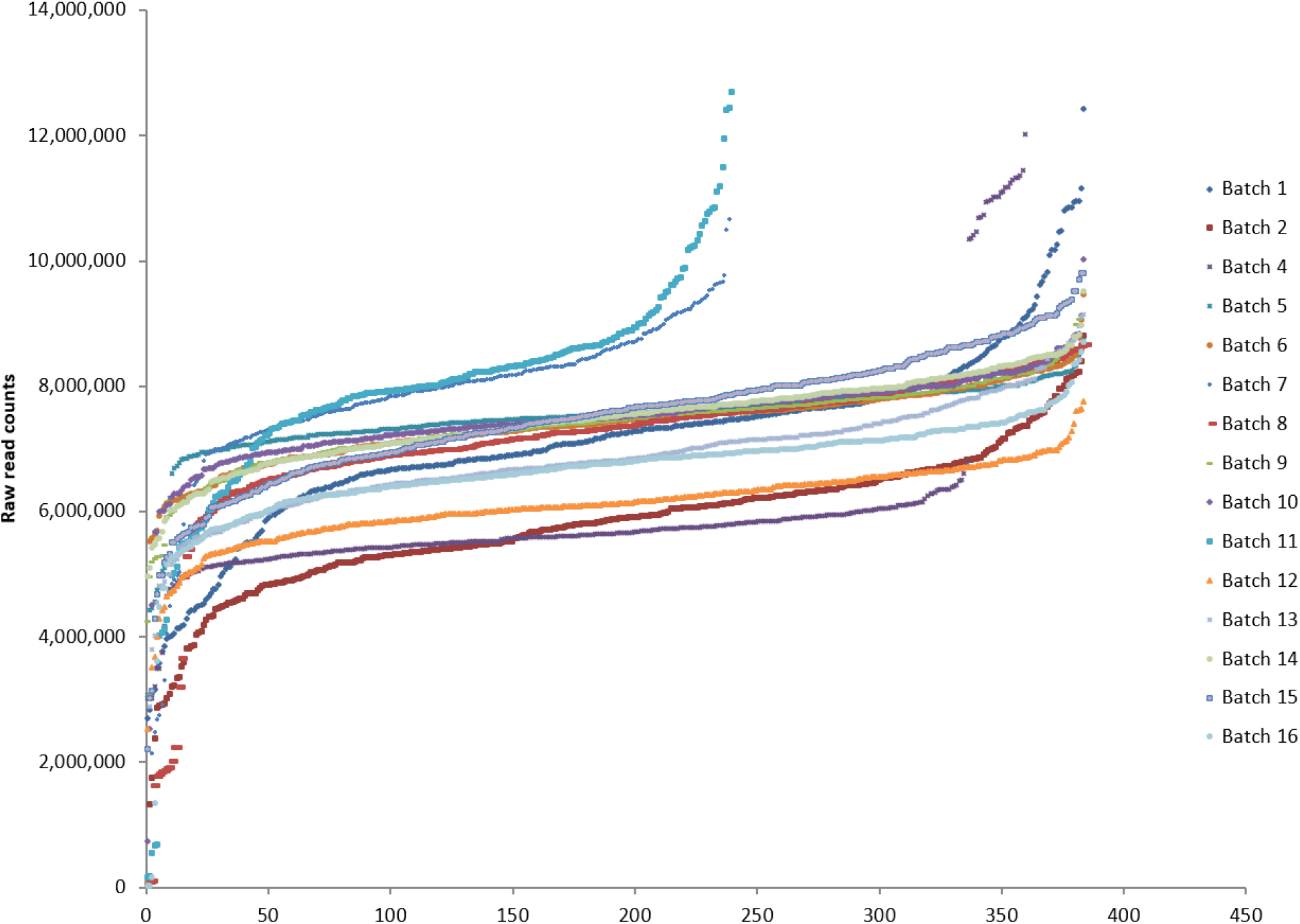
Raw read counts grouped by shipment batch. Raw read counts are on a per-sample basis after demultiplexing FASTQ files with FASTX Barcode Splitter. Each batch represents a set of samples from a given shipment.

**Figure S5.**
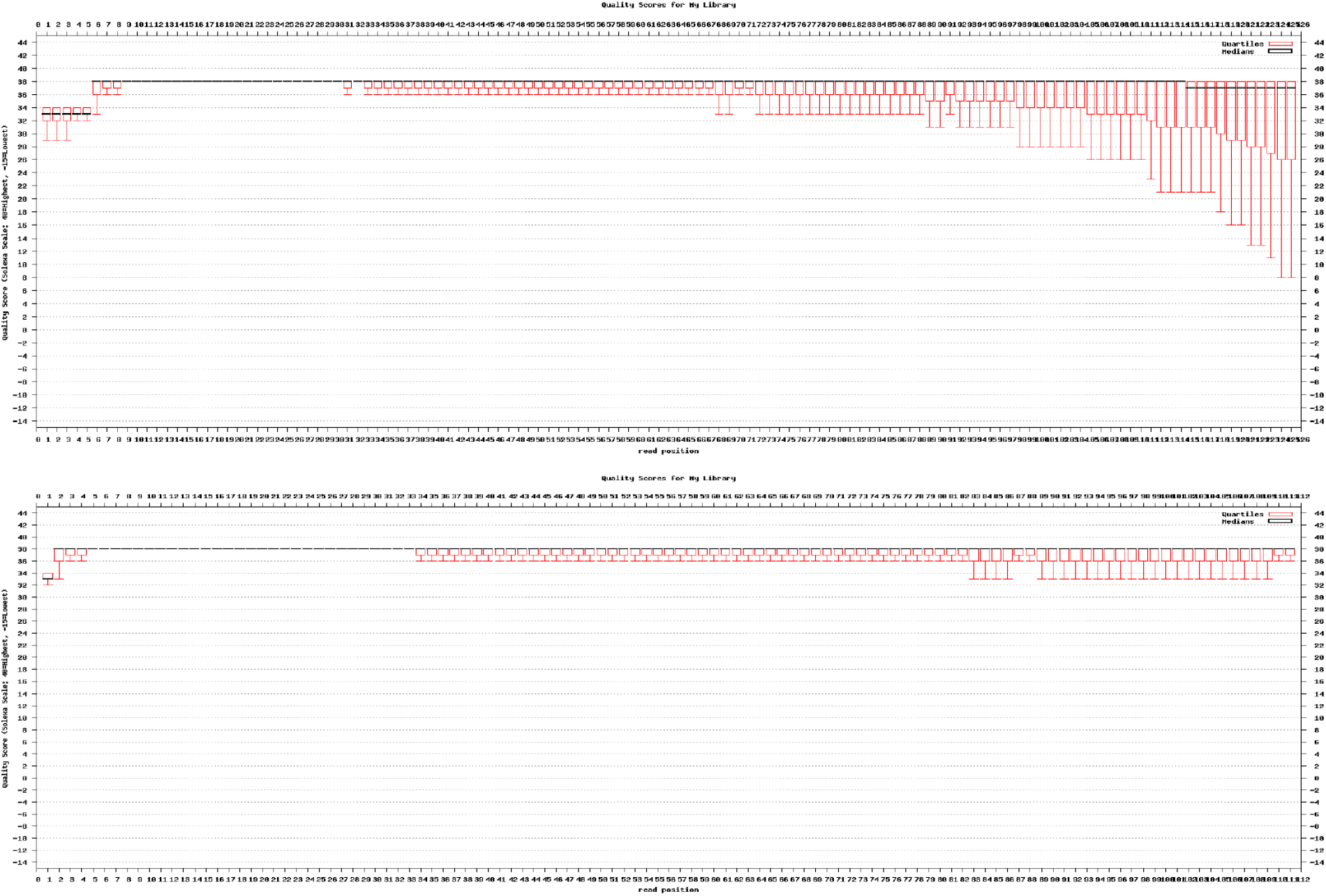
FASTQC results pre- and post-filtering with Cutadapt. FASTQC results are from a single sample from the original set of 96 HS samples prepared in 12-plex and sequenced on the Illumina HiSeq 2500 with 125bp reads.

**Figure S6.**
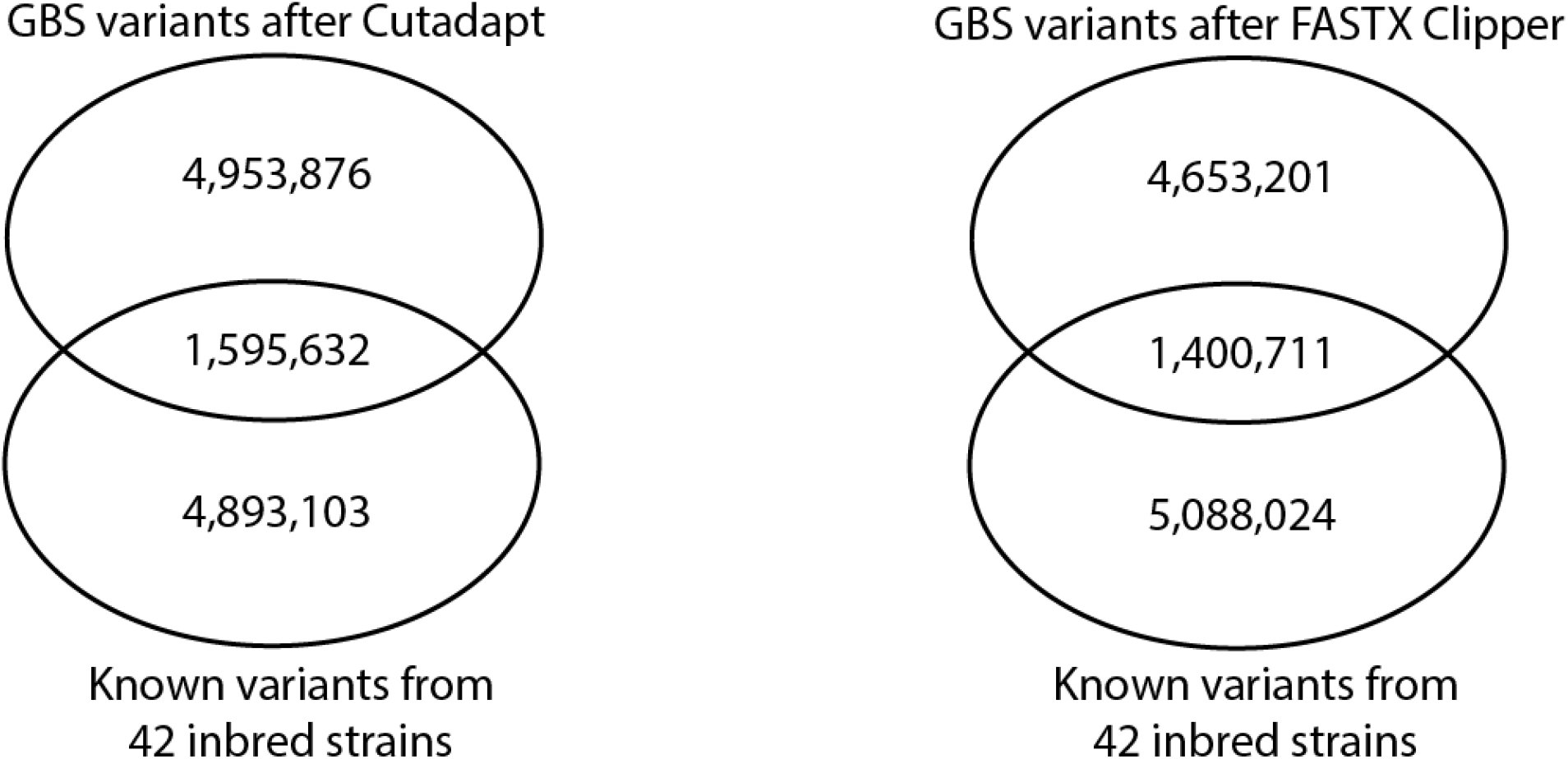
Overlap of called SNPs with known variants after read trimming with FASTX or Cutadapt.

**Figure S7.**
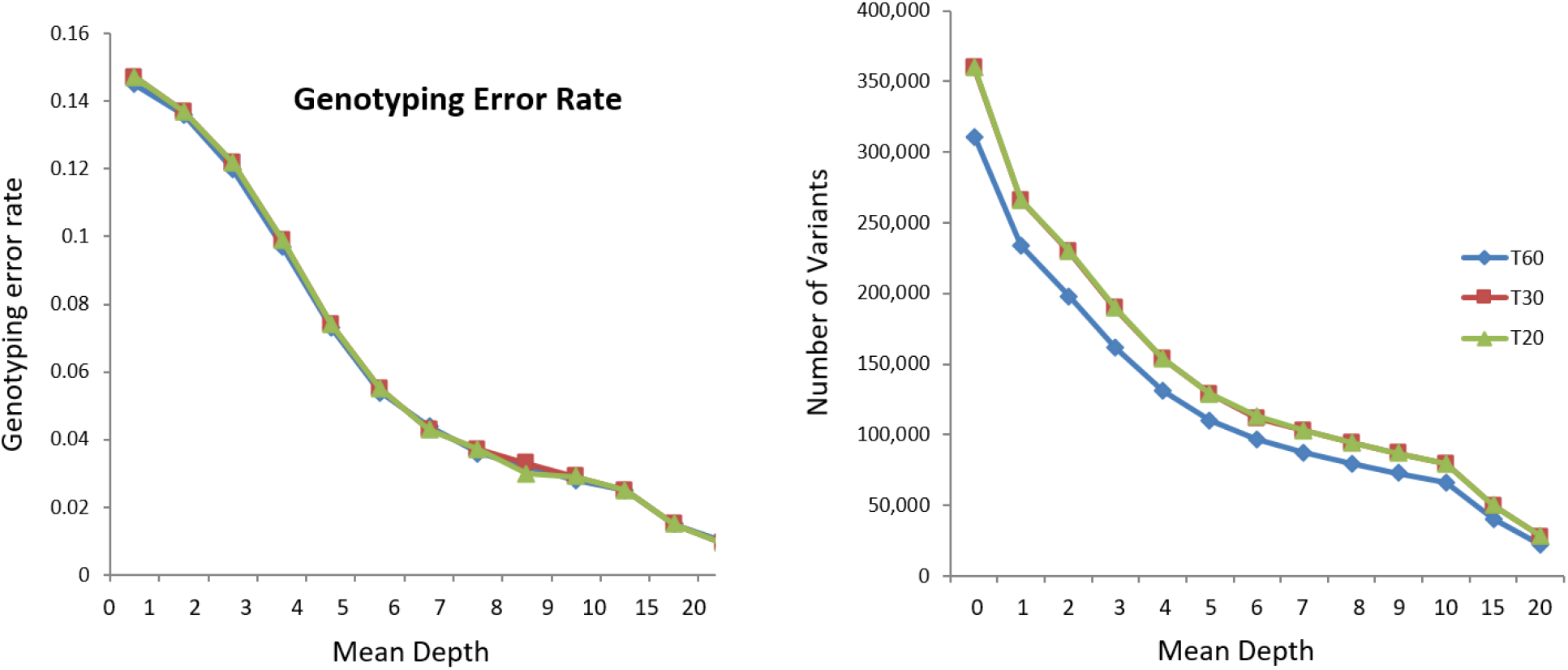
Mapping quality thresholds. Genotyping error rate and number of variants by mean depth per sample per variant site for mapping quality thresholds of 20, 30, and 60.

**Figure S8.**
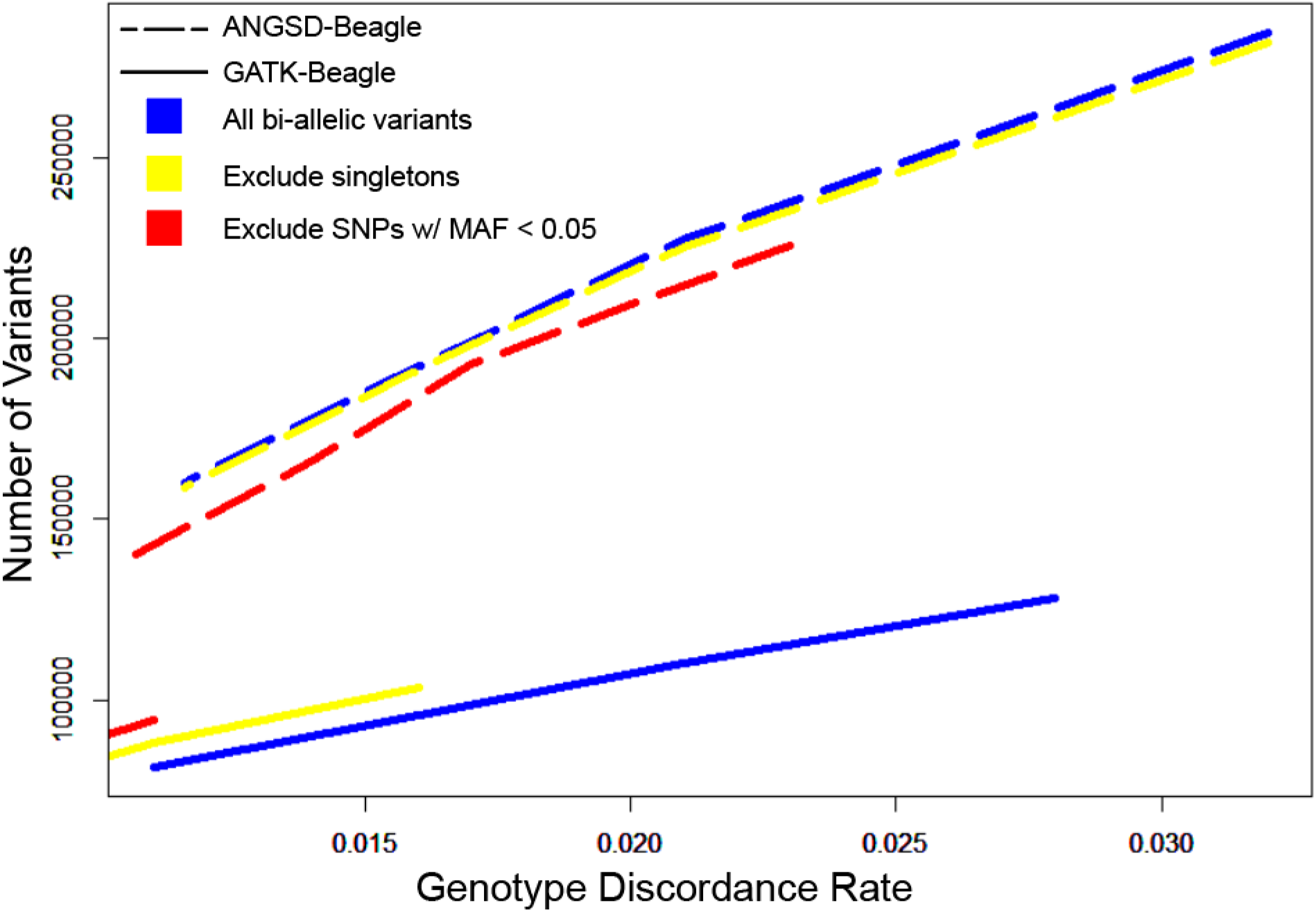
ANGSD-SAMtools vs GATK HaplotypeCaller, filtered calls. The panel compares the number variants called by combination of ANGSD-SAMtools and Beagle or GATK HaplotypeCaller and Beagle at various thresholds of genotype discordance with array data. Calls were made using the 96 HS rats with array data. The x-axis represents the genotype discordance rate thresholds and the y-axis is the number of variants that surpass that threshold for each genotype calling method. Additional filters were applied to the original SNP sets and the plot zooms in on a smaller range of acceptable discordance rates compared to Figure 3. Blue lines represent the unfiltered SNP set. Yellow lines have been filtered for singletons. Red lines have further excluded SNPs with an MAF < 0.05. Each line contains the same number of points.

**Figure S9.**
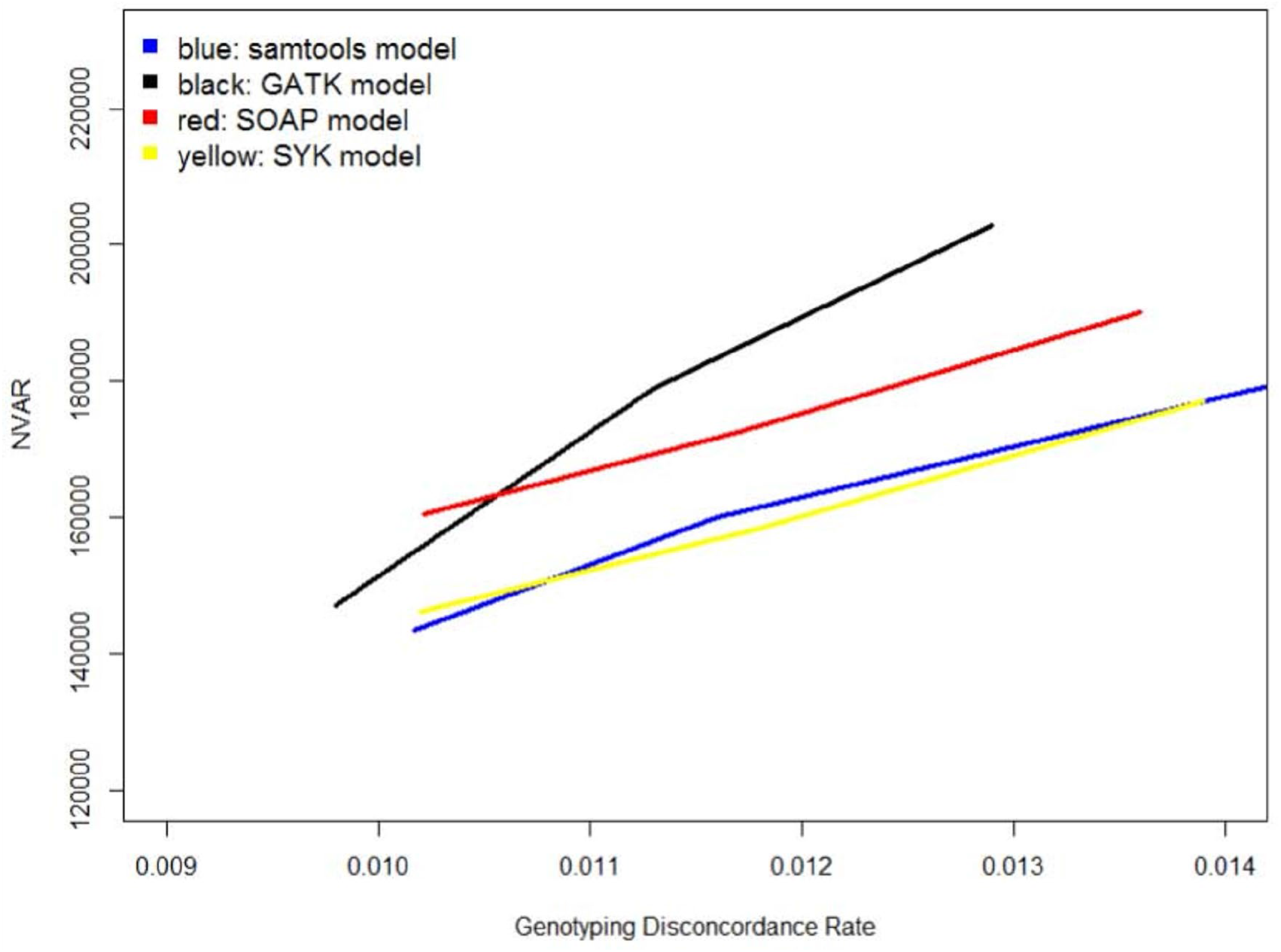
Number of variants by genotype discordance rates for 4 ANGSD genotype likelihood models.

**Figure S10.**
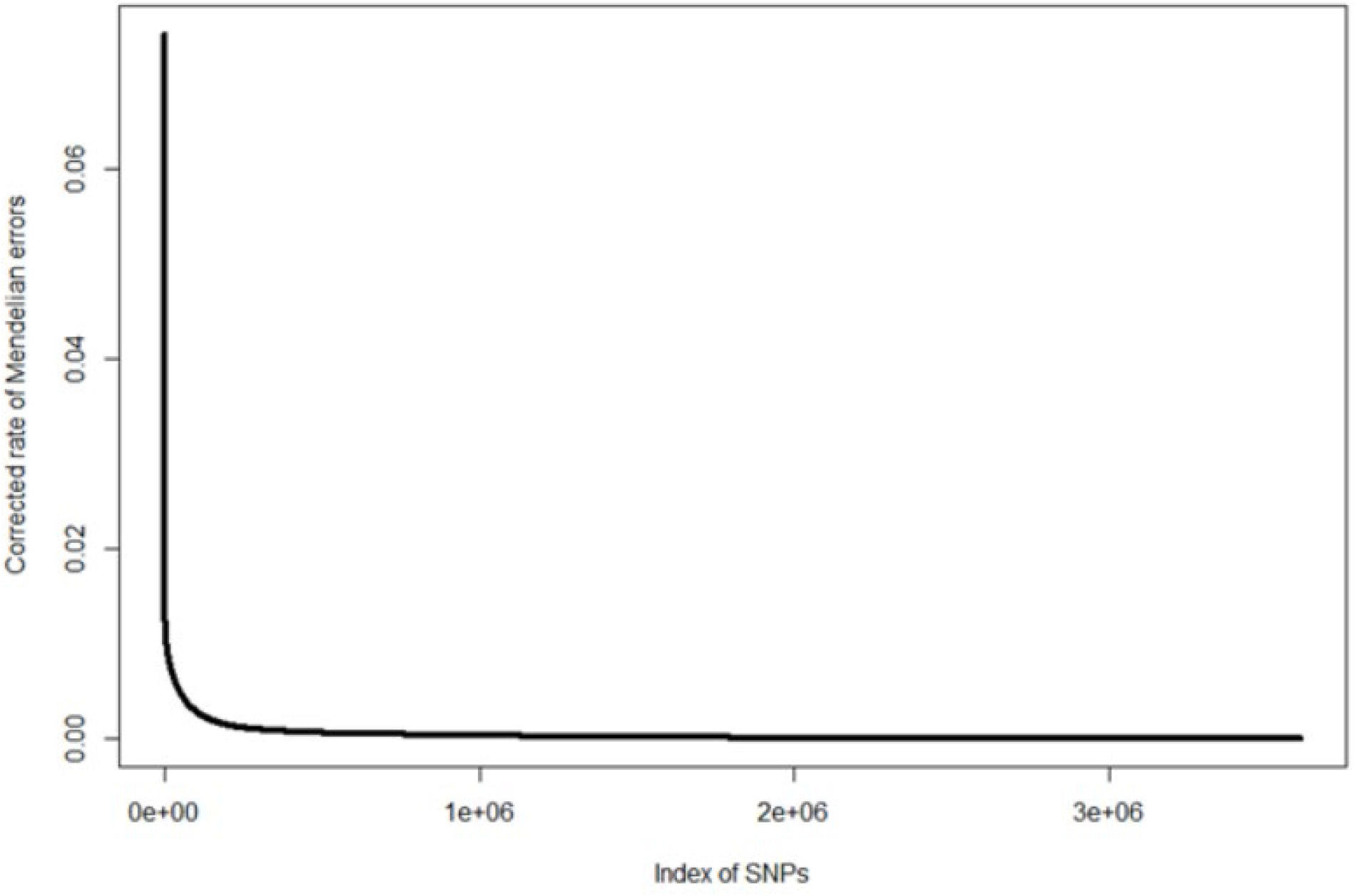
Mendelian error rates. The plot shows the Mendelian error rate for all SNPs. A threshold was set at the inflection point of the curve (~0.005) and all SNPs above that threshold were removed from the data set.

**Figure S11.**
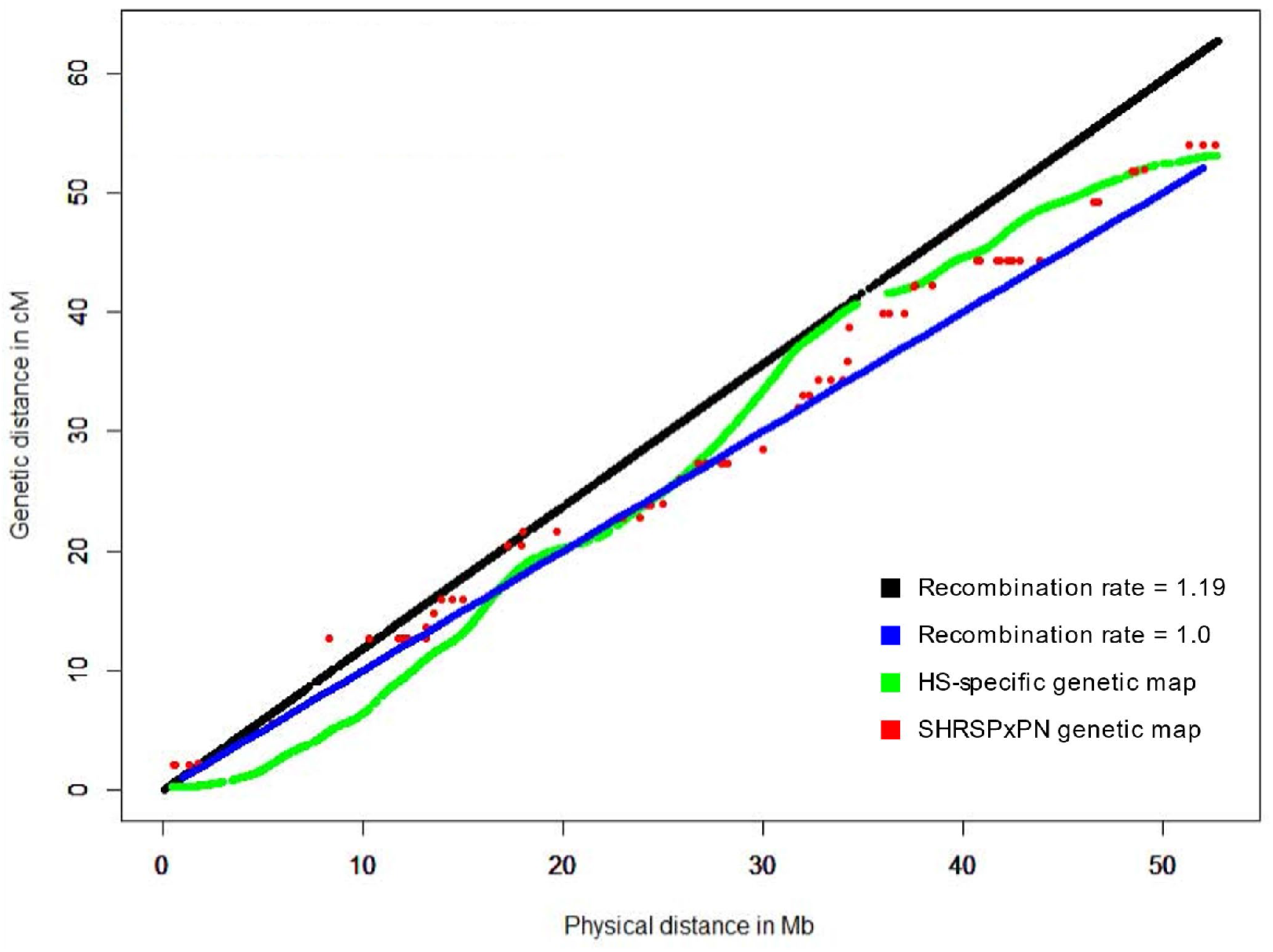
Available rat genetic maps. Plotted physical and genetic distances are for chromosome 12.

**Table S1.**
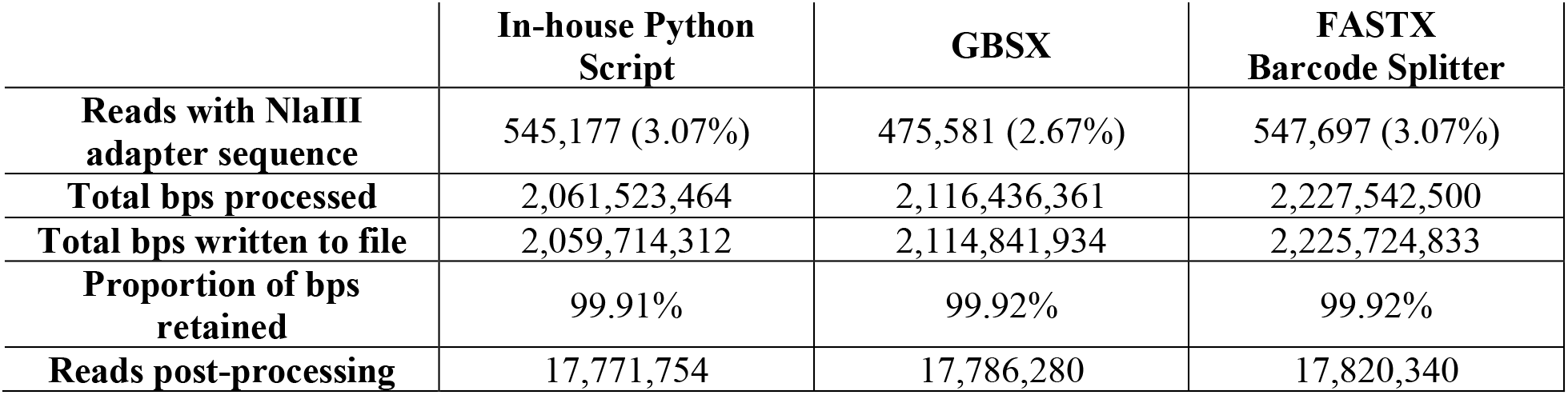
Demultiplexing performance. All methods began with the same number of reads from the original FASTQ. Final read and base pair counts are from after the reads have been trimmed of adapter, barcode, and restriction site sequences, as well as low-quality base pairs (< Q20).

**Table S2.** Comparison of variants calls after filtering with FASTX vs Cutadapt. Data shown comes from the original set of 96 HS samples prepared in 12-plex and sequenced on the Illumina HiSeq 2500. At this early step of pipeline optimization, variants were called utilizing GATK UnifiedGenotyper.

|  | **FASTX Clipper** | **Cutadapt** |
| --- | --- | --- |
| **Number of variants** | 6,075,821 | 6,581,115 |
| **Genotyping call rate** | 0.17 | 0.19 |
| **Mean minor allele count** | 3.96 | 4.25 |
| **Mean minor allele frequency** | 0.15 | 0.15 |
| **Number of singletons** | 433,960 | 548,975 |
| **Number monomorphic sites** | 807,453 | 773,074 |
| **Transition/transversion ratio** | 2.32 | 2.40 |
| **T<sub>I</sub>T<sub>V</sub> ratio for singletons** | 3.23 | 3.40 |
| **Mean variant read depth** | 109.56 | 126.35 |
| **Mean quality score** | 601.79 | 715.56 |

**Table S3.** Variant metrics resulting from reads filtered at different mapping quality thresholds. Data shown comes from the original set of 96 HS samples prepared in 12-plex and sequenced on the Illumina HiSeq 2500. Variants were called utilizing the SAMtools model and the -minMapQ filter in ANGSD. Calls were unfiltered.

|  | MAPQ = 20 | MAPQ = 30 | MAPQ = 45 | MAPQ = 60 | MAPQ = 90 |
| --- | --- | --- | --- | --- | --- |
| **Number of variants** | 372,860 | 372,330 | 363,790 | 316,949 | 233,322 |
| **Genotyping call rate** | 0.64 | 0.64 | 0.64 | 0.61 | 0.75 |
| **Mean minor allele count** | 5.96 | 5.96 | 6.06 | 5.86 | 7.36 |
| **Mean minor allele frequency** | 0.18 | 0.18 | 0.18 | 0.18 | 0.19 |
| **Number of singletons** | 16,781<br>(4.50%) | 16,732<br>(4.49%) | 16,550<br>(4.55%) | 17,352<br>(5.47%) | 11,773<br>(5.05%) |
| **Number of monomorphic sites** | 122,478<br>(32.85%) | 122,188<br>(32.82%) | 116,738<br>(32.09%) | 100,074<br>(31.57%) | 56,179<br>(24.08%) |
| **Transition/transversion ratio** | 1.23 | 1.24 | 1.26 | 1.31 | 1.41 |
| **T<sub>I</sub>T<sub>V</sub> ratio for singletons** | 1.27 | 1.28 | 1.28 | 1.31 | 1.38 |
| **Mean variant read depth** | 157.78 | 157.73 | 159.25 | 152.48 | 188.80 |
| **Mean quality score** | 2,547 | 2,548 | 2,556 | 2,461 | 2,954 |

**Table S4.**
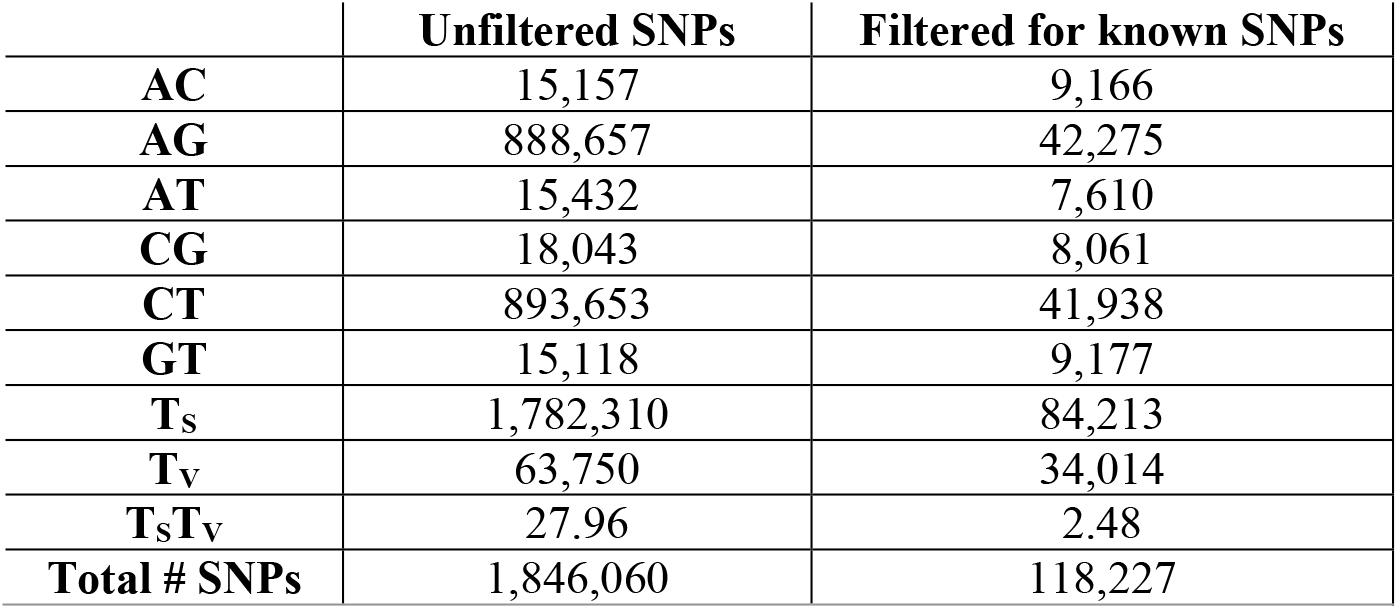
Transition/transversion ratio before and after known sites filtering. The presented data comes from ANGSD-SAMtools/Beagle variant calls for 3,601 HS samples, prior to imputation with IMPUTE2. Known SNPs came from both the 42 inbred genomes from Hermsen et. al 2015 (Hermsen et al. 2015) and the 8 inbred HS founder strains sequenced by the University of Michigan (Ramdas et al. 2019).

|  | Unfiltered SNPs | Filtered for known SNPs |
| --- | --- | --- |
| AC | 15,157 | 9,166 |
| AG | 888,657 | 42,275 |
| AT | 15,432 | 7,610 |
| CG | 18,043 | 8,061 |
| CT | 893,653 | 41,938 |
| GT | 15,118 | 9,177 |
| T<sub>s</sub> | 1,782,310 | 84,213 |
| T<sub>v</sub> | 63,750 | 34,014 |
| T<sub>s</sub>T<sub>v</sub> | 27.96 | 2.48 |
| Total # SNPs | 1,846,060 | 118,227 |

**Table S5.** Imputation accuracy for chromosome 12 across different genetic maps. The number of variants used for the concordance check is dependent on the overlap of the imputed variants with array data for the 96 HS rats with array genotypes. The MAF filter only removes monomorphic sites within the 96 HS rat sample used for the concordance check.

|  | **cM/Mb = 1.00** | **cM/Mb = 1.16** | **SHRSPxPN** | **HS-specific** |
| --- | --- | --- | --- | --- |
| **Number of variants before QC** | 158,452 | 158,452 | 158,452 | 158,452 |
| **Genotyping rate before QC** | 0.94 | 0.92 | 0.92 | 0.92 |
| **Variant removed for missingness > 10%** | 22,217 | 28,959 | 28,356 | 28,858 |
| **Variants removed for MAF < 0.005** | 50,380 | 61,270 | 61,592 | 59,812 |
| **Variants removed for HWE < $1 \times 10^{-10}$** | 53 | 56 | 57 | 56 |
| **Number of variants after QC** | 85,802 | 68,167 | 68,447 | 69,726 |
| **Genotyping rate after QC** | 0.93 | 0.91 | 0.92 | 0.91 |
| **Number of variants in concordance check** | 5,912 | 5,590 | 5,594 | 5,646 |
| **Discordance rate** | 0.095 | 0.011 | 0.011 | 0.010 |

